# SARS-CoV-2 infection leads to Tau pathological signature in neurons

**DOI:** 10.1101/2023.05.17.541098

**Authors:** Cristina Di Primio, Paola Quaranta, Marianna Mignanelli, Giacomo Siano, Matteo Bimbati, Carmen Rita Piazza, Piero Giorgio Spezia, Paola Perrera, Fulvio Basolo, Anello Marcello Poma, Mario Costa, Mauro Pistello, Antonino Cattaneo

**Author notes:** equally contributors (Cristina Di Primio, Paola Quaranta).

## Abstract

**Background:** The coronavirus disease 19 (COVID-19) has represented an issue for global health since its outbreak in March 2020. It is now evident that the SARS-CoV-2 infection results in a wide range of long-term neurological symptoms and is worryingly associated with the aggravation of Alzheimer’s disease. Little is known about the molecular basis of these manifestations.

**Methods:** Several SARS-CoV-2 strain variants were used to infect SH-SY5Y neuroblastoma cells and K18-hACE C57BL/6J mice. The Tau phosphorylation profile and aggregation propensity upon infection were investigated using immunoblot and immunofluorescence on cellular extracts, subcellular fractions, and brain tissue. The viral proteins Spike, Nucleocapsid, and Membrane were overexpressed in SH-SY5Y cells and the direct effect on Tau phosphorylation was checked using immunoblot experiments.

**Results:** Upon infection, Tau is phosphorylated at several pathological epitopes associated with Alzheimer’s disease and other tauopathies. Moreover, this event increases Tau’s propensity to form insoluble aggregates and alters its subcellular localization.

**Conclusions:** Our data support the evidence that SARS-CoV-2 infection in the Central Nervous System triggers downstream effects altering Tau function, eventually leading to the impairment of neuronal function.

## Background

The severe acute respiratory syndrome coronavirus 2 (SARS-CoV-2), which is responsible for coronavirus disease 19 (COVID-19), has been causing a major pandemic with considerable morbidity and mortality since its outbreak in March 2020 (Hu *et al*., 2021; Yong, 2021; Yüce, Filiztekin and Özkaya, 2021).

Despite mainly leading to respiratory disease, increasing studies have reported that this virus can spread to other organs, including the Central Nervous System (CNS). COVID-19 patients present indeed with acute neurological manifestations, such as altered smell and taste, as well as with long-term cognitive symptoms, namely fatigue, headache, attention disorders, dyspnoea, and cognitive impairment (i.e. brain fog) (Ellul *et al*., 2020; Leon *et al*., 2021). This latter condition is defined as “neuro-COVID” and represents a serious concern for global health as it has also been observed in individuals who experienced mild COVID-19 symptoms (Paterson *et al*., 2020; Raveendran, Jayadevan and Sashidharan, 2021). The pathophysiology of this post-viral syndrome is complex as it might involve hypoxia, blood-brain barrier breakage, and, notably, a direct SARS-CoV-2 mediated neuronal tissue invasion and damage. The latter statement is based on the recent confirmation of SARS-CoV-2 neurotropism and the observed loss of grey matter and brain size upon viral infection (Douaud *et al*., 2022). This hypothesis is further corroborated by the brain anatomical and metabolic alterations observed in COVID-19 survivors (Ramani *et al*., 2020; Sun *et al*., 2020; von Weyhern *et al*., 2020; Yong, 2021). Current follow-up studies are beginning to document the feared connection between COVID-19 and neurodegenerative diseases (von Weyhern *et al*., 2020). COVID-19 can exacerbate and accelerate pre-clinical dementia in people with pre-existing Alzheimer’s disease (Ciaccio *et al*., 2021; Emmi *et al*., 2022). At the same time, it leads to a greater propensity for Alzheimer’s disease in patients without pre-existing dementia compared to non-infected individuals, suggesting the association between SARS-CoV-2 and new-onset mental decay (Naughton, Raval and Pasinetti, 2020; Frontera *et al*., 2022).

Despite such reports, to what extent the neuronal effects in patients are a direct consequence of neuronal infection by SARS-CoV-2 or are an indirect effect of the infection of non neuronal cells is still debated (Clijsters *et al*., 2021), the molecular basis for SARS-CoV-2 neuro-infection is yet to be elucidated in healthy neurons or neurodegeneration-primed cells. Within this context, it has been described that COVID-19 patients experiencing neurological symptoms display increased serum levels for brain injury biomarkers, including total Tau and phospho-Tau 181, two hallmarks of Alzheimer’s pathology and other neurodegenerative disorders (Frontera *et al*., 2022; Kokkoris *et al*., 2022).

Here we investigated the interplay between SARS-CoV-2 and the microtubule-associated protein Tau in neurons. Our results indicate that Tau undergoes hyperphosphorylation at pathological epitopes upon infection both *in vitro* and *in vivo*. Moreover, the infection increases Tau’s propensity to accumulate in the insoluble cellular fraction. Both these features are associated with Alzheimer’s disease and other tauopathies, suggesting that SARS-CoV-2 could trigger downstream mechanisms which alter Tau functions.

## Methods

### Cell lines and culture

Human neuroblastoma SH-SY5Y cells (CRL-2266), African green monkey kidney epithelial VeroE6 cells (CRL-1586), and human hepatocyte carcinoma HuH7 cells were purchased from American Type Culture Collection (ATCC, Manassas, VA, USA) and were already available in the lab. VeroE6-TMPRSS2 cells were kindly provided by Dr. Nicola Clementi, Vita-Salute San Raffaele University Hospital, Milan, Italy. SH-SY5Y cells were routinely cultured in Dulbecco’s modified medium: nutrient mixture F12 (DMEM/F12; GIBCO) supplemented with 10% heat-inactivated foetal bovine serum (FBS, EuroClone), 2mM L-glutamine, 100U/mL penicillin and 100 μg/mL Streptomycin; VeroE6, HuH7 and VeroE6-TMPRSS2 cells were maintained in DMEM high glucose medium (Sigma-Aldrich) supplemented with 10% FBS, 2mM L-glutamine, 10U/mL penicillin and 10mg/mL streptomycin. All lines were grown at 37°C with 5% CO_2_.

### Virus isolation and amplification

SARS-CoV-2 manipulation was performed in the biohazard safety level 3 (BLS3) facility of the Virology Unit, Pisa University hospital, Pisa, Italy, and in compliance with the European Committee and the World Health Organization laboratory biosafety guidelines. The SARS-CoV-2 strains used belonged to: B.1 (hCoV-19/Italy/LOM-UniSR10/2021, GISAID Accession ID: EPI_ISL_2544194), B.1.1.7 (hCoV-19/Italy/LOM-UniSR7/2021, GISAID ID: EPI_ISL_1924880), B.1.617.2 (hCoV-19/Italy/TUS-AOUP-FOAN-004/2021 GISAID ID: EPI_ISL_3184308.1). All strains were isolated from nasopharyngeal swab specimens of infected individuals. Briefly, VeroE6 and Vero-TMPRSS2 cells were plated at approximately 80% confluence and infected with 500 μl of the nasal swab buffer diluted in 5 ml of medium. Culture plates were incubated at 37°C, 5% CO_2,_ and shaken every 15 minutes for two hours. At the end, medium supplemented with 5% serum was added and cells cultured until a full cytopathic effect was achieved. Cells were then resuspended in lysis buffer and centrifuged at 900 *g* for 10 min. The supernatants containing virus particles were filtered with a 0.45μM strainer and aliquots were stored at -80°C. Viral stocks were titrated by limited dilution according to the *Reed and Muench* method (REED and MUENCH, 1938) and expressed as Tissue Culture Infectious Dose 50%/ml (TCID50/ml). The titers achieved were: B.1 7×10^8^ TCID50/ml; B.1.1.7 7×10^6^ TCID50/ml; B.1.617.2 5.2×10^6^ TCID50/ml; and B.A.1 1.5×10^6^ TCID50/ml.

### Animals and experimental infection

In vivo procedures were approved by the Ethics Committee for Animal Experimentation of University of Pisa and Italian Ministry of Health (authorization n. 834/2021-PR). Heterozygous *K18-hACE* c57BL/6J mice (strain: 2B6.Cg-Tg(K18-ACE2)2Prlmn/J) were purchased from Charles River Laboratories (Calco, Italy). Animals had *ad libitum* access to standard rodent chow (Mucedola, Milano, Italy) and water under a 12-h light/dark cycle. Briefly, 8-week-old male and female mice were anaesthetised with Ketamine 100 mg/kg and infected by intranasal route with 30 μl of virus SARS-COV-2, B.1 strain, 6×103 TCID50/mice. Animals were monitored daily for the appearance of symptoms and 6 days post infection were sacrificed by cervical dislocation to harvest brains for qRT-PCR, western blot and histology.

### Plasmids and transfection

SARS-CoV-2 Nucleocapsid and Spike plasmids in a pCMV14-3X-Flag backbone were a kind gift from S. Lisi from Bio@SNS laboratory, Scuola Normale Superiore, Italy. The construct encoding SARS-CoV-2 matrix (M) protein was cloned by PCR amplification of M cDNA from the pDONR 207 SARS-CoV-2 M plasmid (Addgene #141274) using a Forward primer inserting a restriction site for EcoRI and a reverse primer bearing a restriction site for BamHI and a Myc tag. proteinM-fwd: 5’-GCT GAA TTC GCA TGG CTG ACT CTA ACG GT-3’; proteinM-rev: 5’-AGC GGA TCC TGC AGA TCC TCT TCA GAG ATG AGT TTC TGC TCC CCC TGC ACC AGC AGG GCG AT-3’. SH-SY5Y were seeded onto 6-well plates (200.000 cells/well) and transfected using the *Lipofectamine 2000* reagent (Thermo Fisher Scientific, Milan, Italy) according to the manufacturer’s instructions. Cells were routinely harvested after 48h of transient expression.

### Cells infection and Western blot analysis

VeroE6, HuH7 and SH-SY5Y cells (2×105 cells/well) were infected with B.1 and B.1.1.7 strains (MOI 0.1). Neuroblastoma cells were additionally challenged with the B.1.617.2 and with B.A.1 variants (MOI 0.01). Following a 2-hour adsorption period, cells were maintained in complete culture medium. 48 hours post-infection, the medium was harvested and stored at -80°C whilst cells were detached with double PBS washes and centrifuged at 900 *g* for 10 min. Total protein extraction was carried out in RIPA buffer (Millipore, USA) for infected cells and in aTris-HCl 20 mM pH 8,1, NaCl 20 mM, glycerol 10%, NP-40 1%, EDTA 10 mM buffer for transfected cells. Both reagents were supplemented with protease and phosphatase inhibitors (Roche). The extracts were centrifuged at 16,000 rcf for 15 min at 4°C to collect the supernatants, which were quantified with Bradford method before loading buffer resuspension. Western Blotting experiments on viral particles were carried out on the collected culture media concentrated with *Vivaspin 6 MWCO 30 kDa membranes* (Sigma Aldrich). Samples were then reduced and denatured by boiling in 4X Laemmli sample buffer at 95°C for 5 minutes. Triton-X 100 fractionation was performed as previously reported in *Siano et al., 2019* (Siano, Caiazza, Ollà, *et al*., 2019). Briefly, infected cell pellets were resuspended in 1% Triton-X 100 in PBS lysis buffer supplemented with protease and phosphatase inhibitors. Lysates were subsequently centrifuged at 16000 x g for 15 minutes at 4°C. Pellets (Triton X-100 insoluble fractions) were further dissolved in 1% SDS; 1% Triton-X in PBS lysis buffer, whereas supernatants (Triton X-100 soluble fraction) were directly processed. Brains excised from infected and control *K18-hACE* mice were homogenised in Ripa buffer added with proteinase and phosphatase inhibitors and incubated overnight at 4°C. The following day, extracted proteins were quantified by Bradford method and processed for western blot analysis as follows. Equal amounts of protein preparations were separated by 10% Tris-Glycine SDS-PAGE SDS-PAGE (Bio-Rad) and then transferred onto a nitrocellulose membrane (Amersham Biosciences). Membranes were blocked with EveryBlot Blocking buffer solution (Bio-Rad) for 5 minutes at room temperature according to the manufacturer’s instruction and hybridized with the primary antibody in the same blocking buffer overnight at 4°C. The following day, incubation with HRP-conjugated antibodies diluted in blocking solution was carried out at room temperature for 1 hour. Membrane washing was performed 3 times for 5 minutes in Tris-buffered saline with 0,1% Tween20 (TBST). Blots were developed with SuperSignal™ West Femto Maximum Sensitivity Substrate (ThermoFisher Scientific) and acquired by ChemiDoc™ Imaging Systems (Bio-Rad). Primary antibodies: Rabbit SARS-CoV-2 Spike S1 antibody [HL6], 1.1000 (GTX635654, GeneTex, Irvine, CA); mouse SARS-CoV / SARS-CoV-2 Spike S2 antibody [1A9], 1.1000 (GTX632604, GeneTex, Irvine, CA); rabbit SARS-CoV-2 Nucleocapsid antibody, 1.1000 (GTX135357, GeneTexm Irvine, CA); mouse anti-Tau (Tau5), 1.1000 (ab80579, Abcam) mouse anti-tau (Tau13) 1.1000 (sc-21796 Santa Cruz Biotechnology, Dallas, TX); rabbit anti-pTau (Ser262),1.500 (OPA1-03142, ThermoFisher Scientific); mouse anti p-Tau (Ser396), 1.500 (#9632, Cell Signaling technology, Danvers, MA); mouse anti p-Tau Thr231, 1.500 (# MN1040, ThermoFisher Scientific); mouse anti-GAPDH, 1.10000 (10R-G109a, Fitzgerald Industries international, Acton, MA; rabbit anti-β-tubulin, 1.1000 (#2146, Cell Signaling Technology); rabbit anti-histone H2B, 1.1000 (sc-515808, Santa Cruz Biotechnology, Dallas, GTX); rabbit anti-actin, 1.5000 (A300-485A, Bethyl Laboratories); HRP-Conjugated anti-Myc-Tag (9B11) (#2040 Cell Signaling Technology). Secondary antibodies: goat anti-mouse IgG-HRP, 1.0000 (sc-516102, Santa Cruz Biotechnology, Dallas,TX); mouse anti-rabbit IgG-HRP, 1.10000(sc-2357, Santa Cruz Biotechnology, Dallas, TX)

### Immunofluorescence

SH-SY5Y (10.000 cells/well) were seeded onto 8 well-chamber microscope slides (Nunc Lab Tech, ThermoFisher Scientific) and infected with the SARS-CoV-2 B.1 strain (MOI 0.1) as described above. After a 24-h adsorption period, cells were washed from the viral inoculum 3 times in Phosphate Buffered Saline (PBS) and fixed with 100% ice-cold methanol for 15 minutes.

Samples were then permeabilized in Triton 0,1% in PBS for 5 minutes and blocked in a 1% BSA-Tween 0,1% in PBS solution for 30 minutes at room temperature. Hybridization with primary antibodies diluted in blocking solution was performed overnight at 4°C. The following day, cells were permeabilized and blocked again for 5 minutes and incubated with Alexa Fluor-Conjugated secondary antibodies diluted in blocking solution for 1 hour at room temperature. Nuclei were stained with DAPI (Sigma Aldrich). Brain tissues were fixed in buffered formalin solution and stored at 4°C overnight. The day after, they were included in OCT solution (Bio-Optica, Milan, Italy) and cut in 50 μm coronal slices with a freezing microtome. Free-floating sections were then blocked in 10% Normal Goat Serum (NGS) - 0.3% Triton in PBS 1X for 2 hours at room temperature and then hybridised with primary antibodies overnight at 4°C. On the following day, slices were incubated with the secondary antibody for 2 hours at room temperature. Both primary and secondary antibodies were diluted in a 1% NGS-0.1% Triton-PBS solution. Primary antibodies: anti-S IgG rabbit monoclonal antibody (40592-V05H, Sino Biological); rabbit SARS-CoV-2 Nucleocapsid antibody, 1.1000 (GTX135357, GeneTex Irvine, CA); anti-α-tubulin IgG mouse monoclonal antibody (T5168, Merck); mouse anti-tau (Tau13), 1.500 (sc-21796 Santa Cruz Biotechnology, Dallas, TX); Secondary antibodies (1.200 dilution) :goat anti-rabbit IgG Alexa488-labeled monoclonal antibody; donkey anti-mouse IgG Alexa568-labeled monoclonal antibody); goat anti-mouse IgG Alexa546-labeled monoclonal antibody goat anti-mouse IgG Alexa633-labeled monoclonal antibody (ThermoFisher Scientific); DAPI, 1.10000 (28718-90-3, Sigma Aldrich); TOTO-3 iodide, 1.5000 (T3604, ThermoFisher Scientific). Images were acquired on a Zeiss Laser Scanning (LSM) 880 confocal microscope (Carl Zeiss, Jena, Germany) supplied with GaAsP (Gallium:Arsenide:Phosphide) detectors. Samples were viewed with a 63X Apochromat oil immersion (1.4 NA) DIC objective. Whole-cell images were acquired with a z-stack series of usually ten slices with 0.5μm intervals and summed up with the z-projection tool from Fiji.

### SARS-CoV-2 Real-Time PCR

Weighted brains were stored in trizol reagent and homogenised at 4°C, the RNA was extracted by Chomczynski protocol and quantified and assessed for purity by nanodrop. qRT-PCR was performed to quantify the SARS-COV2 viral load present in the tissue. SARS-CoV-2 RNA relative amounts detected for each experimental condition as a cycle threshold (Ct) value were compared, with a mean Ct value determined for the positive infection control. The purified RNA was then used to perform the synthesis of first-strand complementary DNA, using the One-Step TB Green® PrimeScript™ RT-PCR Kit II (Takara) for RT-PCR. Real-time PCR, using the SYBR Green dye– based PCR amplification and detection method, was performed to detect the complementary DNA. We used the forward primer SF(CTCATCACGTAGTCGCAACAGTTC) the reverse primer SR (CAAGCTGGTTCAATCCTGTCAAGCA) for Spike protein, and the forward primer AF (CTCCATCCTGGCCTCACTGT) and the reverse primer AR (GAGGGGCCGGACTCATCGT) for actin. The PCR conditions were: 42°C for 2 minutes, 45 cycles of 95°C for 10 seconds, annealing at 95°C for 5 seconds and elongation at 62°C for 30 seconds, followed by a final elongation at 72°C for 10 minutes. RT-PCR was performed using the ABI-PRISM 7900HT Fast Real Time instrument (Applied Biosystems) and optical-grade 96-well plates. Samples were run in duplicate, with a total volume of 20μL.

### Statistical analysis

In western blot assays, differences between means were analysed by the non-parametric Mann-Whitney test (for n=2 independent experimental groups) or Kruskal-Wallis test followed by post-hoc Mann-Whitney (for n>2 independent experimental groups). Fold changes were calculated with respect to control groups, to which a theoretical mean of 1 was attributed. All results are shown as mean (bars) ± SEM (whiskers) from at least three independent experiments. Levels of significance are depicted as * for p < 0.05, ** for p < 0.01, *** for p < 0.001, n.s. non-significant. Origin 9.0 (OriginLab, Northampton, MA) software was used to perform statistical analysis and plotting.

## Results

### SARS-CoV-2 is associated with Tau in human neuron-like cells

To investigate SARS-CoV-2 dynamics and its downstream effects in neurons, we infected SH-SY5Y cells, a neuroblastoma line widely used as an in *vitro* model of neurodegenerative disorders and competent for productive infection with SARS-CoV-2 (Gordon, Amini and White, 2013; Yin, Baillie and Vetter, 2016; Siano, Caiazza, Ollà, *et al*., 2019; Siano, Caiazza, Varisco, *et al*., 2019; Slanzi *et al*., 2020; Benedetti *et al*., 2021; Bielarz *et al*., 2021; Torices *et al*., 2021; Bartolomeo *et al*., 2022).

Three clinical strains belonging to the following variants: B.1, B.1.1.7, B.1.617.2 have been used, and the infection verified by immunoblot assays and confocal microscopy imaging 48 hours post-infection. We detected Spike (S) and Nucleocapsid (N) viral proteins in both infected neuroblastoma and control VeroE6 and HuH7 cells, which are routinely used for viral amplification and isolation (*Figure 1A*).

**Figure 1.**
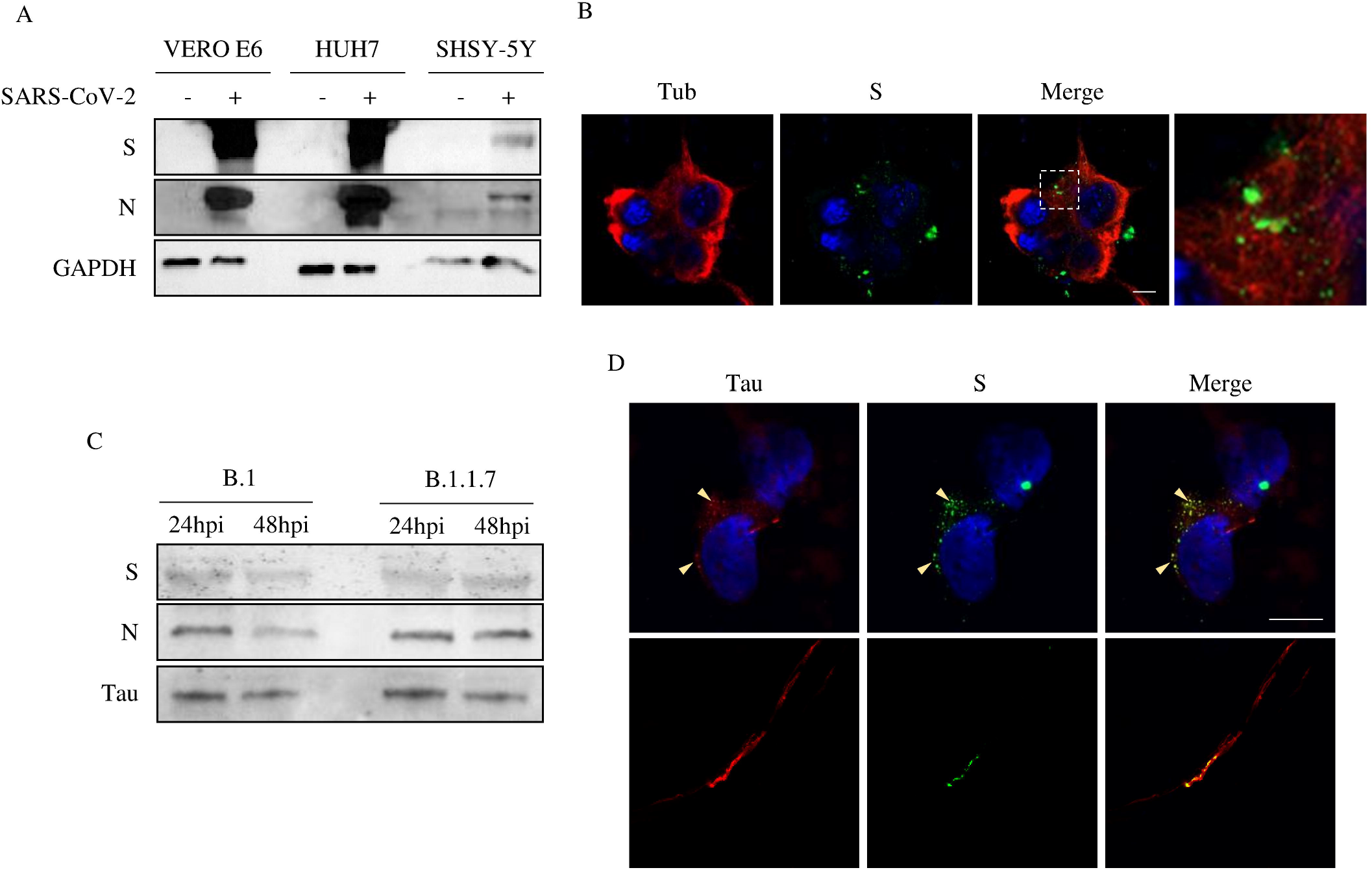
SARS-Cov2 infects human neuron-like cells. Viruses detected in infected cells. (A) Viral proteins detected in cellular extracts of infected and control cells by immunoblot. Spike (S), Nucleocapsid (N). (B) Super resolution imaging of viral particles detected in infected SH-SY5Y cells 48 hours post-infection. SARS-CoV-2 spike (green), Tubulin (red), DAPI (blue). (C) Immunoblot of virions released 24 and 48 hpi in the supernatant of SH-SY5Y cells infected with B.1 or B.1.1.7 strains. (D) Super resolution imaging of infected cells. SARS-CoV-2 S (green), Tau (red), DAPI (blue). Scale bar:10 μm

As shown by the Tubulin and S staining in F*igure 1B*, infected cells displayed an intact microtubule network; although only partially, a few viral particles appeared aligned along the microtubule tracks. Immunoblot assays on collected cell media further showed that SH-SY5Y cells underwent productive infection, as new viral particles were generated and released into the medium. Unexpectedly, the viral progeny contained, or was associated with, the microtubule-associated protein Tau *(Figure 1C)*.

We further investigated the interplay between viral particles and Tau by super-resolution imaging of infected cells. As shown in *Figure 1D*, viral puncta co-localized with the Tau signal. Remarkably, Tau formed spots facing viral signals.

These data suggest that SARS-CoV-2, like other CoVs (Wen *et al*., 2020), might hijack the neuronal cytoskeleton during the viral life cycle in neurons. In particular, the involvement of Tau in the viral life cycle in both infected cells and newly-formed virions suggests possible downstream implications for neuronal homeostasis and highlights a novel aspect of SARS-CoV-2 molecular pathology possibly based on cell-to-cell Tau spreading.

### SARS-CoV-2 leads to Tau hyperphosphorylation and aggregation

It has been reported that SARS-CoV-2 induces a global phosphorylation signaling as a primary host response to the infection (Bouhaddou *et al*., 2020; Stukalov *et al*., 2021; Xia, Wang and Zheng, 2021; Chatterjee and Thakur, 2022; Liu *et al*., 2022; Boytz *et al*., 2023; Maginnis, 2023). *Surjit et al*. (Surjit *et al*., 2004) previously demonstrated that SARS-CoV N protein significantly up-regulates p38 mitogen-activated protein kinase (MAPK) cascade, whose activation is involved in actin remodeling, neuroinflammation, and, notably, Tau hyperphosphorylation (Mizutani *et al*., 2004). Recent studies have confirmed that SARS-CoV-2 activates p38 activity in infected cells likewise (Grimes and Grimes, 2020; Goel *et al*., 2021).

Therefore, we assessed the Tau phosphorylation profile at epitopes considered pathological hallmarks of numerous neurodegenerative disorders (Avila, 2009; Alonso *et al*., 2010). By immunoblot experiments we found a significant increase in Tau phosphorylation at Ser262 (p262) and Ser396 (p396); however, the signal from Tau p231 was unaltered (*Figure 2A; Additional file 1*). Of note, all tested viral strains lead to the same phosphorylation profile, suggesting that their differences in transmissibility have no impact on downstream Tau modifications.

**Figure 2.**
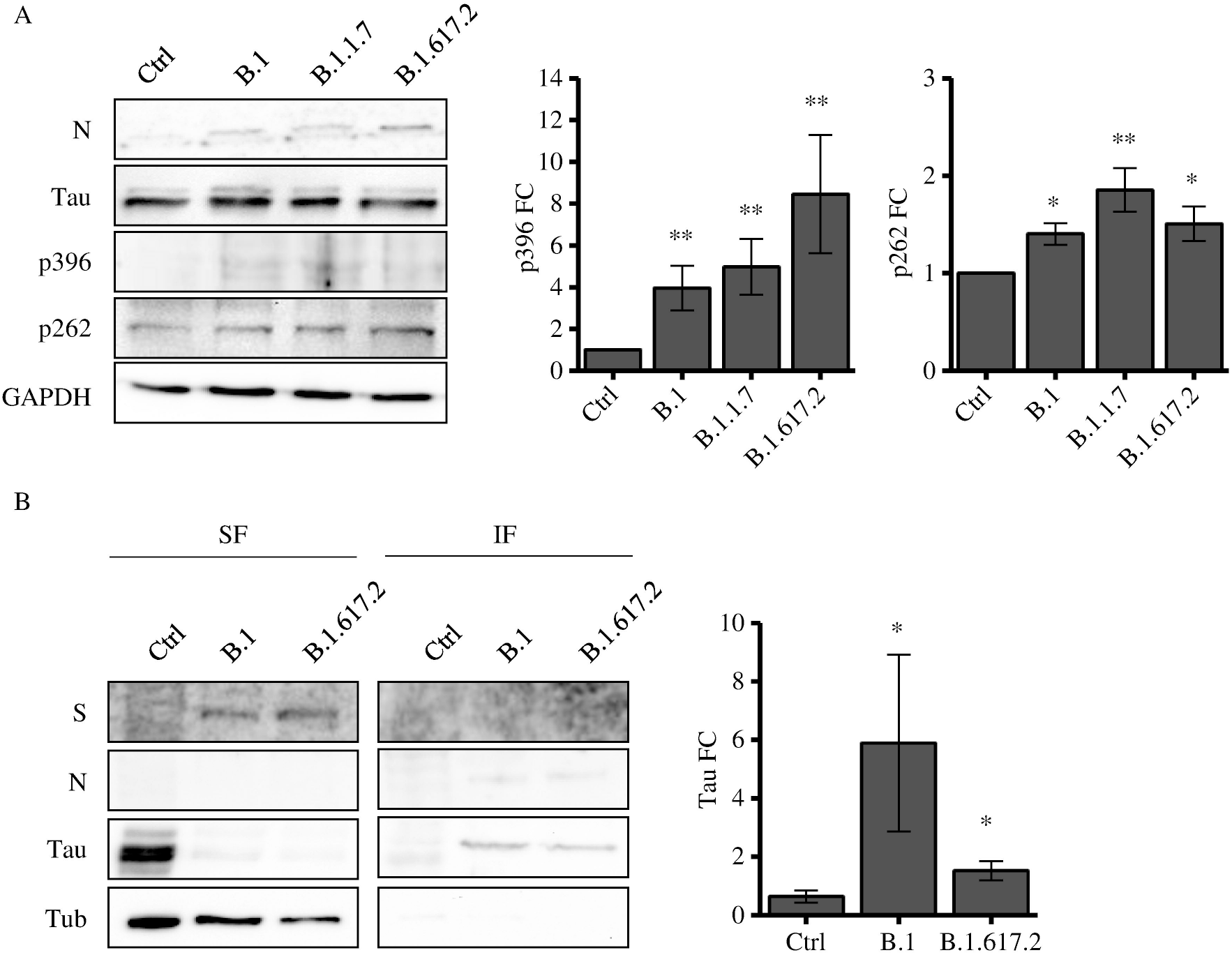
Tau is phosphorylated and accumulates in the insoluble fraction upon SARS-CoV-2 infection. (A) Immunoblot analysis of SH-SY5Y cells not infected (N.I.) and infected with SARS-CoV-2 strains and relative quantification (n=5/9) Kruskal-Wallis non-parametric ANOVA followed by Mann-Whitney post-hoc comparison; *p<0.05. (B) Immunoblot of soluble and insoluble cellular fractions after detergent fractionation of cells infected with B.1 and B.1.1.7 strains and control cells. (C) Quantification of Tau in the soluble fraction. (D) Quantification of Tau in the insoluble fraction. Kruskal-Wallis ANOVA and Mann-Whitney test; *p<0.05.

It is well known that Tau hyperphosphorylation disrupts its interaction with microtubules, thereby leading to its mislocalization and increasing its propensity to form insoluble aggregates (Johnson and Stoothoff, 2004; Avila *et al*., 2022). We checked the presence of Tau in the soluble and insoluble fractions of infected cells, and we found a robust accumulation of Tau in the insoluble fraction (*Figure 2 B,C,D*). Again, the formation of insoluble species was significantly induced by all the tested viral strains similarly. Remarkably, the S protein could be observed in the soluble fraction whereas protein N was enriched in the insoluble fraction with Tau, suggesting a putative interplay between these proteins, which might account for Tau pathological alterations.

To further investigate this aspect and gain insights into the molecular mechanism underlying SARS-CoV-2-dependent Tau phosphorylation, we examined whether the virus itself triggered these alterations. Hence, we dissected the viral components and tested their separate ability to impact Tau phosphorylation.

To this aim, we transfected neuroblastoma cells with plasmids expressing SARS-CoV-2 N and Membrane (M). As shown in figure 3, either N or M expression induced a significant increase in Tau p262, while p231 and p396 were not modulated. On the contrary, S expression did not alter Tau phosphorylation at these residues (*Additional file 2*). These data suggest a direct role for both SARS-CoV-2 N and M in altering Tau function, which might potentially trigger downstream events leading to neuro-COVID.

**Figure 3.**
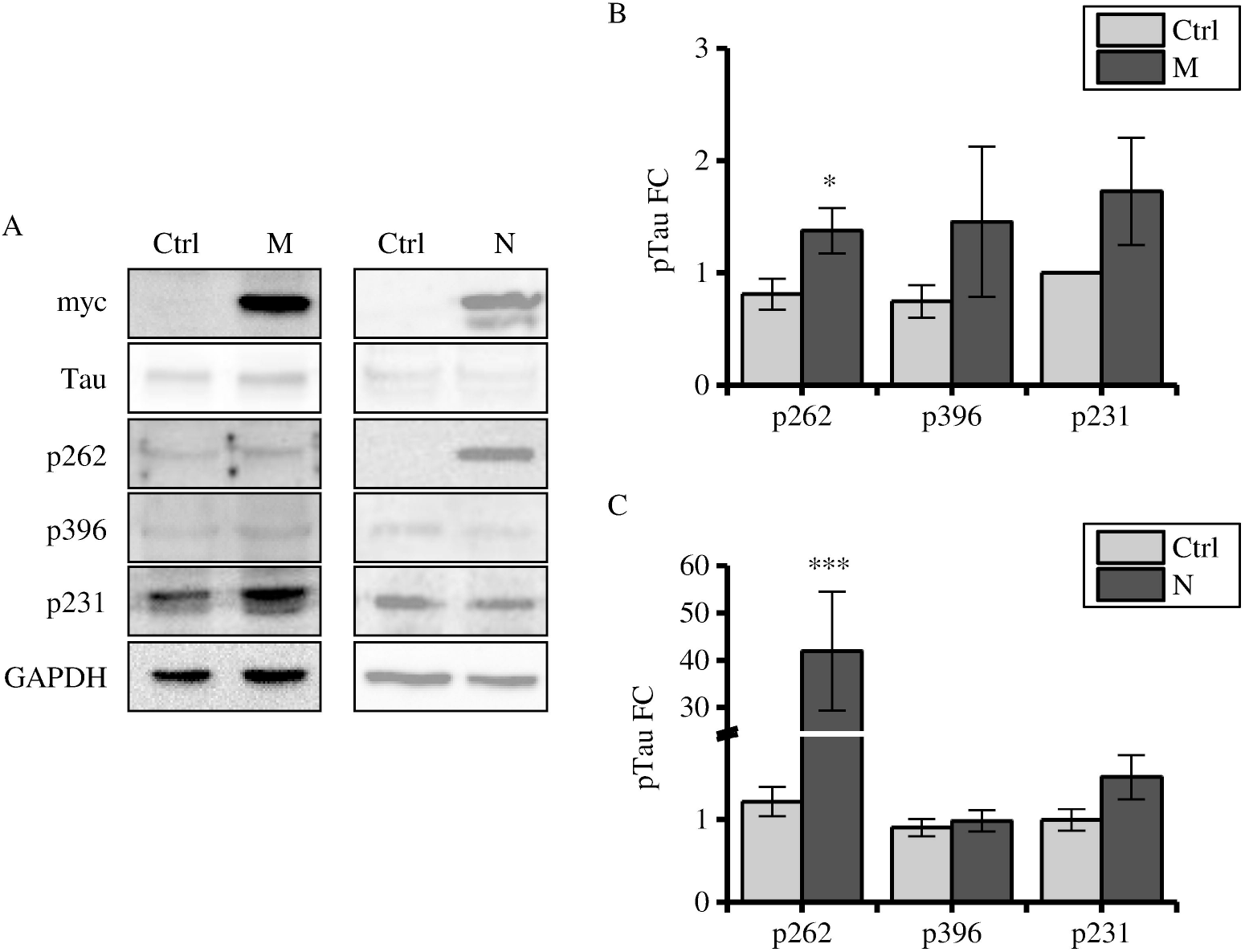
N and M lead to Tau phosphorylation at S262. (A) Immunoblot analyses of cells expressing N and control cells. Relative quantification of p231, p396, AT8, p262. Mann-Whitney; ***=p<0.001. (B) Immunoblot analyses of cells expressing M and control cells. Relative quantification of p231, p396, p262. Mann-Whitney; *=p<0.05.

Taken together, our findings suggest that SARS-CoV-2 infection of human sustains Tau hyperphosphorylation at several pathological epitopes, an effect possibly mediated by its integral proteins N and M, which eventually leads to its mislocalization and aggregation in neuroblastoma cells.

### SARS-CoV-2 leads to Tau phosphorylation and mislocalization in the mouse brain

Given our findings in neuron-like cells, we assessed Tau modulation *in vivo* by challenging a transgenic mouse line expressing human ACE2 (K18-*hACE2* model (Winkler *et al*., 2020)) with the SARS-CoV-2 B.1 viral strain. As previously shown by *Winkler et al*., infected mice developed severe respiratory syndrome, which eventually led to premature death within about 7 days.

By immunoblot analysis on brain tissues from mice sacrificed 6 days after infection (Methods), we observed that SARS-CoV-2 infection caused a significant increase of Tau p262 and p396 (*Figure 4A*).

**Figure 4.**
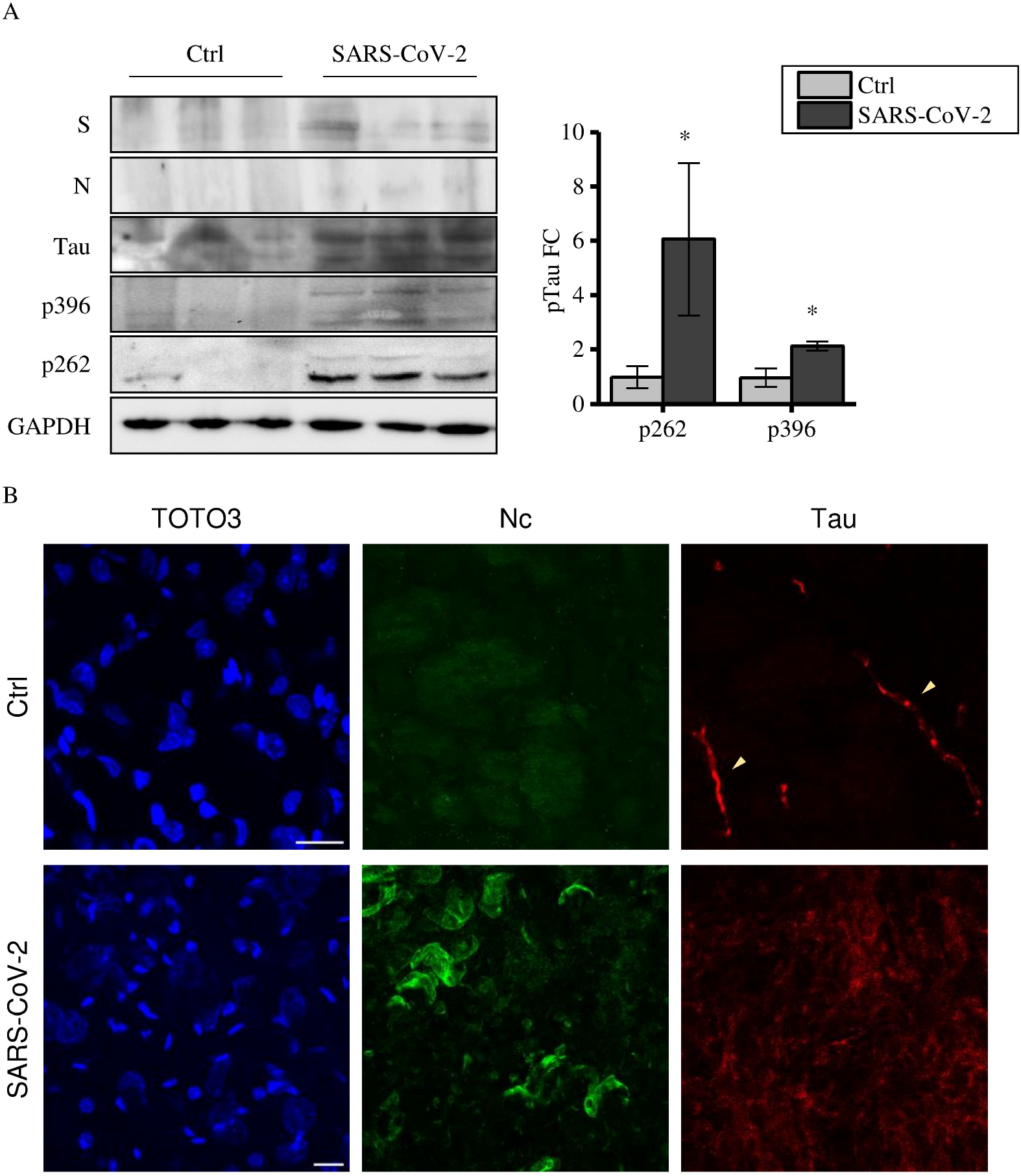
Tau is hyperphosphorylated and mislocalized in the infected mouse brain. (A) Immunoblot assay of brain lysates from K18-hACE2 mice after SARS-CoV-2 infection and relative quantification; non parametric Mann-Whitney: *p-value<0,05; ** p-value<0,01. (B) Confocal images of brain cross sections from K18-hACE2 mice either uninfected (N.I.) or infected (SARS-CoV-2) with B.1. strain. Nuclei stained with TOTO3 (blue); N (green); Tau (red). Scale bar:20 μm

To understand whether this modulation altered Tau subcellular distribution and function, we assessed its localization in neurons. As shown in figure 4B, brain slices from control animals displayed Tau enriched at neurites decorating the neuronal cytoskeleton, as usually observed. However, Tau appeared widespread in the cytoplasm after viral infection. This data supports the evidence that SARS-CoV-2 infection in the CNS might result in symptoms commonly associated with neurodegenerative disorders due to its involvement in Tau post-translational modification and, eventually, its mislocalization.

## Discussion

New SARS-CoV-2 variants are continuously emerging causing new waves of infection and major worldwide burden. Although most COVID-19 patients primarily develop respiratory signs, they may experience neurological symptoms with cognitive and psychiatric impairments, which may persist after the acute phase of infection (Paterson *et al*., 2020; Leon *et al*., 2021; Raveendran, Jayadevan and Sashidharan, 2021). This new illness is of great concern as it might affect mental and social well-being by interfering with daily life.

As the virus keeps mutating and spreading worldwide, it is a matter of necessity and urgency to provide a molecular mechanism underlying COVID-19-related long-term neurological complications. Filling this gap would eventually contribute to finding strategies for treatment and prevention. This work provides new insights by exploring viral downstream effects in neurons.

Given that SARS-CoV-2 neurotropism has been widely proven to be comparable to other Coronaviruses, we exploited SH-SY5Y cells as a human neuronal-like cellular model to investigate the molecular biology of the viral-host interplay (Chen *et al*., 2020; Ramani *et al*., 2020; Wang *et al*., 2021; Mesci *et al*., 2022).

In these cells, the viral replication level is compared to VeroE6 and HuH7 cells, which are the gold standard platforms to amplify and isolate this virus. Although their infection rate was lower than that of reference cells, SH-SY5Y cells exhibited active viral replication, and the viral progeny they released was associated with the neuronal protein Tau. This event was not surprising since several enveloped virions from different taxa have been described to package host cell proteins into or onto their surface (Me and Tremblay, 2005). However, this finding needs further investigation due to its potential long-term complications. A common mechanism for the progression of several neurodegenerative disorders is indeed the transcellular transmission of pathological proteins and the subsequent misfolding of their wild-type counterparts in recipient neurons (Peng, Trojanowski and Lee, 2020). Within this context, a well-studied pathway is the spreading of hyperphosphorylated Tau aggregates in the brains of Alzheimer’s disease patients (Saman *et al*., 2012; Wang *et al*., 2017).

It is widely recognized that most neurotropic viruses, including *betacoronaviruses* like SARS and MERS, as well as HIV-1, HSV, and ZIKV, hijack the cytoskeleton network to fulfil a successful infection. The fine interplay between viral and host cytoskeletal elements is relevant for the pathogen spreading across the cytoplasm towards the ERGIC compartment or the nucleus (Lehmann, Nikolic and Piguet, 2011; Wen *et al*., 2020; Nie *et al*., 2021). In this work, we observed that only few viral particles co-localize with tubulin on the microtubules network but there is a strong association with Tau, suggesting that the SARS-CoV-2 neurotropic mechanism employs the neuronal cytoskeleton. In addition, we found that the virus interplay with Tau results in increased phosphorylation at Ser262 and Ser396 residues both *in vitro* and *in vivo*. These epitopes are widely described as hallmarks of Alzheimer’s disease and other tauopathies (Avila, 2009; Alonso *et al*., 2010). It is known that hyperphosphorylated Tau displays decreased interaction with microtubules, and certain residues such as Ser262 are the major sites inhibiting this binding efficiency (Drewes *et al*., 1995). This event could cause the formation of insoluble aggregates, which are linked with neuron toxicity and are considered the main trigger of tauopathies (Alonso *et al*., 2001). In line with these data, we demonstrated that SARS-CoV-2 infection is associated with Tau overspreading and mislocalization in the mouse brain. This observation is also in agreement with high-resolution imaging data from SARS-CoV-2-infected human brain organoids^8^, showing that it is associated with altered distribution of Tau from axons to soma. Moreover, we detected a significant increase of Tau in the insoluble fraction of infected cells, which indicates that SARS-CoV-2 affects not only the phospho-Tau profile, but also its biochemical properties. These findings suggest that SARS-CoV-2 infection in neurons causes Tau pathological changes, which could eventually contribute to neuro-COVID symptoms by impairing neuronal homeostasis.

Previous work has shown the association between SARS-CoV and SARS-CoV-2 infections and the increased activity of the p38 mitogen-activated protein kinase (MAPK) pathway, which is involved in actin remodelling and Tau phosphorylation (Mizutani *et al*., 2004; Surjit *et al*., 2004; Grimes and Grimes, 2020). Future research should address which kinases responsible for Tau phosphorylation are over-activated upon viral entry in neurons. Of note, the Ser396 residue is more susceptible to proline-rich kinases as glycogen synthase kinase 3-beta(Li and Paudel, 2006). MAPKs such as the extracellular signal-regulated kinases 1 and 2, c-Jun amino-terminal kinases, and p38 γ and δ isoforms are involved in neurodegeneration as well (Bouhaddou *et al*., 2020; Falcicchia *et al*., 2020).

It is still unclear whether Tau aberrant phosphorylation profile and its consequent aggregation are caused by the virus itself upon infection or/and by an indirect cellular response. In this regard, we observed that the expression of SARS-CoV-2 Membrane and Nucleocapsid proteins is sufficient to increase Tau phosphorylation, indicating that Tau post-translational modulation could be at least partially modulated by a direct function of these viral proteins, as already described for their SARS-CoV orthologues (Mizutani *et al*., 2004; Surjit *et al*., 2004). By making microtubules less stable, Tau phosphorylation could represent a cell defensive strategy to detach the virus from microtubules, preventing its transport to the ERGIC compartment, i.e., the assembly and budding site of newly-synthesised virions. Of note, pTau profile is not altered by the expression of Spike, which is subject to a higher mutation rate than N and M (Abavisani *et al*., 2022), which potentially highlights these two viral proteins as a druggable target to counteract COVID-dependent neuropathology.

## Conclusions

We demonstrated that SARS-CoV-2 infection in neuronal cells triggers aberrant Tau phosphorylation at several pathological epitopes which are associated with Alzheimer’s disease and other tauopathies. By altering Tau properties, this event eventually leads to its aggregation and the impairment of neuronal function as a consequence of the displacement of host cytoskeletal components. Our data open up a new molecular mechanism underlying post-COVID neurological manifestations and acknowledge the potential scale of the disease long-term course. Although vaccination is currently the primary strategy for the management of the SARS-CoV-2 pandemic, patients who already experienced COVID-19 must not be side-lined. To limit long-term consequences, it is relevant for clinicians to be aware of the downstream pathophysiological aberrations when dealing with neuro-COVID cases.

## List of abbreviations

SARS-CoV-2: Severe Acute Respiratory Syndrome Coronavirus 2
COVID-19: Coronavirus disease 19
CNS: Central Nervous System
CoVs: Coronaviruses
N: Nucleocapsid
M: Membrane
S: Spike
pTau: phospho-Tau

## Declarations

### Ethics approval and consent to participate

The current study was approved by Ministero della Salute, Governo Italiano, authorization n° 834/2021-PR, art. 31 del D.lgs. 26/2014 and performed accordingly.

### Consent for publication

All authors approve the content of this manuscript and agree to publication

### Availability of data and material

Data and material are available upon request

### Authors’ contributions

Conceptualization: C.D.P. and P.Q. Methodology: C.D.P., P.Q., M.M., M.P., F.B. and M.C. Investigation: C.D.P., P.Q., M.M., G.S., M.B., C.R.P. and P.P. Supervision: C.D.P., M.P, A.C. and M.C. Writing: C.D.P., M.M., P.Q., A.C. and all authors commented and approved the final version.

## Acknowledgements

The authors are grateful to Alessandro Viegi, Simonetta Lisi and Elena Novelli for technical support.

**Additional files 1**.

.**Tiff**

**Tau phosphorylation at the p-231 epitope**.

Immunoblot analysis of SH-SY5Y cells infected with SARS-CoV-2 B.1, B.1.1.7, B.1.617.2 strains; n= 5/7 experiments. Kruskal-Wallis non-parametric ANOVA: n.s.

**Additional files 2**.

.**Tiff**

**S does not modulate Tau phosphorylation**.

Immunoblot analyses of cells expressing S and control cells.

## References

Abavisani, M. et al. (2022) ‘Mutations in SARS [CoV [2 structural proteins[: a global analysis’, pp. 1–19. doi: 10.1186/s12985-022-01951-7.

Alonso, A. et al. (2001) ‘Hyperphosphorylation induces self-assembly of tau into tangles of paired helical filaments/straight filaments.’, Proceedings of the National Academy of Sciences of the United States of America, 98(12), pp. 6923–6928. doi: 10.1073/pnas.121119298.

Alonso, A. D. et al. (2010) ‘Phosphorylation of tau at Thr212, Thr231, and Ser262 combined causes neurodegeneration.’, The Journal of biological chemistry, 285(40), pp. 30851–30860. doi: 10.1074/jbc.M110.110957.

Avila, J. (2009) ‘The tau code’, 1(July), pp. 1–5. doi: 10.3389/neuro.24.001.2009.

Avila, S. et al. (2022) ‘Role of Tau Protein in Both Physiological and Pathological Conditions’, pp. 361–384.

Bartolomeo, C. S. et al. (2022) ‘SARS-CoV-2 infection and replication kinetics in different human cell types: The role of autophagy, cellular metabolism and ACE2 expression.’, Life sciences. Netherlands, 308, p. 120930. doi: 10.1016/j.lfs.2022.120930.

Benedetti, F. et al. (2021) ‘Comparison of SARS-CoV-2 Receptors Expression in Primary Endothelial Cells and Retinoic Acid-Differentiated Human Neuronal Cells.’, Viruses. Switzerland, 13(11). doi: 10.3390/v13112193.

Bielarz, V. et al. (2021) ‘Susceptibility of neuroblastoma and glioblastoma cell lines to SARS-CoV-2 infection.’, Brain research. Netherlands, 1758, p. 147344. doi: 10.1016/j.brainres.2021.147344.

Bouhaddou, M. et al. (2020) ‘The Global Phosphorylation Landscape of SARS-CoV-2 Infection.’, Cell. United States, 182(3), pp. 685–712.e19. doi: 10.1016/j.cell.2020.06.034.

Boytz, R. et al. (2023) ‘Anti-SARS-CoV-2 activity of targeted kinase inhibitors: Repurposing clinically available drugs for COVID-19 therapy.’, Journal of medical virology. United States, 95(1), p. e28157. doi: 10.1002/jmv.28157.

Chatterjee, B. and Thakur, S. S. (2022) ‘SARS-CoV-2 Infection Triggers Phosphorylation: Potential Target for Anti-COVID-19 Therapeutics.’, Frontiers in immunology. Switzerland, 13, p. 829474. doi: 10.3389/fimmu.2022.829474.

Chen, R. et al. (2020) ‘The Spatial and Cell-Type Distribution of SARS-CoV-2 Receptor ACE2 in the Human and Mouse Brains.’, Frontiers in neurology. Switzerland, 11, p. 573095. doi: 10.3389/fneur.2020.573095.

Ciaccio, M. et al. (2021) ‘COVID-19 and Alzheimer’s Disease.’, Brain sciences. Switzerland, 11(3). doi: 10.3390/brainsci11030305.

Clijsters, M. et al. (2021) ‘Article Visualizing in deceased COVID-19 patients how SARS-CoV-2 attacks the respiratory and olfactory mucosae but spares the olfactory bulb ll ll Visualizing in deceased COVID-19 patients how SARS-CoV-2 attacks the respiratory and olfactory mucosae but spares the olfactory bulb’, Cell. Elsevier Inc., 184(24), p. 5932–5949.e15. doi: 10.1016/j.cell.2021.10.027.

Douaud, G. et al. (2022) ‘SARS-CoV-2 is associated with changes in brain structure in UK Biobank’. Springer US, 604(August 2021). doi: 10.1038/s41586-022-04569-5.

Drewes, G. et al. (1995) ‘Microtubule-associated protein/microtubule affinity-regulating kinase (p110mark). A novel protein kinase that regulates tau-microtubule interactions and dynamic instability by phosphorylation at the Alzheimer-specific site serine 262.’, The Journal of biological chemistry. United States, 270(13), p. 7679–7688. doi: 10.1074/jbc.270.13.7679.

Ellul, M. A. et al. (2020) ‘Since January 2020 Elsevier has created a COVID-19 resource centre with free information in English and Mandarin on the novel coronavirus COVID-research that is available on the COVID-19 resource centre - including this for unrestricted research re-use and analyses in any form or by any means with acknowledgement of the original source. These permissions are Rapid Review Neurological associations of COVID-19’, (January).

Emmi, A. et al. (2022) ‘Smell deficits in COVID-19 and possible links with Parkinson’s disease.’, International review of neurobiology. United States, 165, p. 91–102. doi: 10.1016/bs.irn.2022.08.001.

Falcicchia, C. et al. (2020) ‘Involvement of p38 MAPK in Synaptic Function and Dysfunction.’, International journal of molecular sciences, 21(16). doi: 10.3390/ijms21165624.

Frontera, J. A. et al. (2022) ‘Comparison of serum neurodegenerative biomarkers among hospitalized COVID-19 patients versus non-COVID subjects with normal cognition, mild cognitive impairment, or Alzheimer’s dementia.’, Alzheimer’s & dementiaJ: the journal of the Alzheimer’s Association. United States, 18(5), p. 899–910. doi: 10.1002/alz.12556.

Goel, S. et al. (2021) ‘SARS-CoV-2 Switches “ on “ MAPK and NF κ B Signaling via the Reduction of Nuclear DUSP1 and DUSP5 Expression’, 12(April), p. 1–11. doi: 10.3389/fphar.2021.631879.

Gordon, J., Amini, S. and White, M. K. (2013) ‘General overview of neuronal cell culture.’, Methods in molecular biology (Clifton, N.J.). United States, p. 1–8. doi: 10.1007/978-1-62703-640-5_1.

Grimes, J. M. and Grimes, K. V (2020) ‘Since January 2020 Elsevier has created a COVID-19 resource centre with free information in English and Mandarin on the novel coronavirus COVID-19. The COVID-19 resource centre is hosted on Elsevier Connect, the company ‘ s public news and information website. Elsevier hereby grants permission to make all its COVID-19-related research that is available on the COVID-19 resource centre - including this research content - immediately available in PubMed Central and other publicly funded repositories, such as the WHO COVID database with rights for unrestricted research re-use and analyses in any form or by any means with acknowledgement of the original source. These permissions are granted for free by Elsevier for as long as the COVID-19 resource centre remains active. p38 MAPK inhibition[: A promising therapeutic approach for COVID-19’, (January).

Hu, B. et al. (2021) ‘Characteristics of SARS-CoV-2 and COVID-19.’, Nature reviews. Microbiology, 19(3), p. 141–154. doi: 10.1038/s41579-020-00459-7.

Johnson, G. V. W. and Stoothoff, W. H. (2004) ‘Tau phosphorylation in neuronal cell function and dysfunction’. doi: 10.1242/jcs.01558.

Kokkoris, S. et al. (2022) ‘Serum inflammatory and brain injury biomarkers in COVID-19 patients admitted to intensive care unit: A pilot study.’, eNeurologicalSci. Netherlands, 29, p. 100434. doi: 10.1016/j.ensci.2022.100434.

Lehmann, M., Nikolic, D. S. and Piguet, V. (2011) ‘How HIV-1 Takes Advantage of the Cytoskeleton during Replication and Cell-to-Cell Transmission’, p. 1757–1776. doi: 10.3390/v3091757.

Leon, S. L. et al. (2021) ‘More than 50 long [term effects of COVID [19[: a systematic review and meta [analysis Middle East respiratory syndrome’, Scientific Reports. Nature Publishing Group UK, p. 1–12. doi: 10.1038/s41598-021-95565-8.

Li, T. and Paudel, H. K. (2006) ‘Glycogen synthase kinase 3beta phosphorylates Alzheimer’s disease-specific Ser396 of microtubule-associated protein tau by a sequential mechanism.’, Biochemistry. United States, 45(10), p. 3125–3133. doi: 10.1021/bi051634r.

Liu, X. et al. (2022) ‘SARS-CoV-2 spike protein-induced cell fusion activates the cGAS-STING pathway and the interferon response.’, Science signaling. United States, 15(729), p. eabg8744. doi: 10.1126/scisignal.abg8744.

Maginnis, M. S. (2023) ‘β-arrestins and G protein-coupled receptor kinases in viral entry: A graphical review.’, Cellular signalling. England, 102, p. 110558. doi: 10.1016/j.cellsig.2022.110558.

Me, S. and Tremblay, M. J. (2005) ‘MINIREVIEW Plunder and Stowaways[: Incorporation of Cellular Proteins by Enveloped Viruses Re’, 79(11), p. 6577–6587. doi: 10.1128/JVI.79.11.6577.

Mesci, P. et al. (2022) ‘SARS-CoV-2 infects human brain organoids causing cell death and loss of synapses that can be rescued by treatment with Sofosbuvir.’, PLoS biology, 20(11), p. e3001845. doi: 10.1371/journal.pbio.3001845.

Mizutani, T. et al. (2004) ‘Phosphorylation of p38 MAPK and its downstream targets in SARS coronavirus-infected cells.’, Biochemical and biophysical research communications, 319(4), p. 1228–1234. doi: 10.1016/j.bbrc.2004.05.107.

Naughton, S. X., Raval, U. and Pasinetti, G. M. (2020) ‘Potential Novel Role of COVID-19 in Alzheimer’s Disease and Preventative Mitigation Strategies.’, Journal of Alzheimer’s diseaseJ: JAD. Netherlands, 76(1), p. 21–25. doi: 10.3233/JAD-200537.

Nie, Y. et al. (2021) ‘Rearrangement of Actin Cytoskeleton by Zika Virus Infection Facilitates Blood – Testis Barrier Hyperpermeability’, Virologica Sinica. Springer Singapore, 36(4), p. 692–705. doi: 10.1007/s12250-020-00343-x.

Paterson, R. W. et al. (2020) ‘The emerging spectrum of COVID-19 neurology[: clinical, radiological and laboratory findings’, p. 3104–3120. doi: 10.1093/brain/awaa240.

Peng, C., Trojanowski, J. Q. and Lee, V. M.-Y. (2020) ‘Protein transmission in neurodegenerative disease.’, Nature reviews. Neurology. England, 16(4), p. 199–212. doi: 10.1038/s41582-020-0333-7.

Ramani, A. et al. (2020) ‘SARS-CoV-2 targets neurons of 3 D human brain organoids’, p. 1–14. doi: 10.15252/embj.2020106230.

Raveendran, A. V, Jayadevan, R. and Sashidharan, S. (2021) ‘Long COVID: An overview.’, Diabetes & metabolic syndrome, 15(3), p. 869–875. doi: 10.1016/j.dsx.2021.04.007.

Reed, L. J. and H. (1938) ‘A SIMPLE METHOD OF ESTIMATING FIFTY PER CENT ENDPOINTS12’, American Journal of Epidemiology, 27(3), p. 493–497. doi: 10.1093/oxfordjournals.aje.a118408.

Saman, Sudad et al. (2012) ‘Exosome-associated tau is secreted in tauopathy models and is selectively phosphorylated in cerebrospinal fluid in early Alzheimer disease.’, The Journal of biological chemistry. United States, 287(6), p. 3842–3849. doi: 10.1074/jbc.M111.277061.

Siano, G., Caiazza, M. C., Ollà, I., et al. (2019) ‘Identification of an ERK Inhibitor as a Therapeutic Drug Against Tau Aggregation in a New Cell-Based Assay.’, Frontiers in cellular neuroscience. Switzerland, 13, p. 386. doi: 10.3389/fncel.2019.00386.

Siano, G., Caiazza, M. C., Varisco, M., et al. (2019) ‘Modulation of Tau Subcellular Localization as a Tool to Investigate the Expression of Disease-related Genes’, (December), p. 1–9. doi: 10.3791/59988.

Slanzi, A. et al. (2020) ‘In vitro Models of Neurodegenerative Diseases.’, Frontiers in cell and developmental biology. Switzerland, 8, p. 328. doi: 10.3389/fcell.2020.00328.

Stukalov, A. et al. (2021) ‘Multilevel proteomics reveals host perturbations by SARS-CoV-2 and SARS-CoV.’, Nature. England, 594(7862), p. 246–252. doi: 10.1038/s41586-021-03493-4.

Sun, S.-H. et al. (2020) ‘A Mouse Model of SARS-CoV-2 Infection and Pathogenesis.’, Cell host & microbe, 28(1), p. 124–133.e4. doi: 10.1016/j.chom.2020.05.020.

Surjit, M. et al. (2004) ‘The SARS coronavirus nucleocapsid protein induces actin reorganization and apoptosis in COS-1 cells in the absence of growth factors.’, The Biochemical journal. England, 383(Pt 1), p. 13–18. doi: 10.1042/BJ20040984.

Torices, S. et al. (2021) ‘Expression of SARS-CoV-2-related receptors in cells of the neurovascular unit[: implications for HIV-1 infection’. Journal of Neuroinflammation, 2, p. 1–16.

Wang, C. et al. (2021) ‘ApoE-Isoform-Dependent SARS-CoV-2 Neurotropism and Cellular Response.’, Cell stem cell. United States, 28(2), p. 331–342.e5. doi: 10.1016/j.stem.2020.12.018.

Wang, Y. et al. (2017) ‘The release and trans-synaptic transmission of Tau via exosomes.’, Molecular neurodegeneration. England, 12(1), p. 5. doi: 10.1186/s13024-016-0143-y.

Wen, Z. et al. (2020) ‘Cytoskeleton-a crucial key in host cell for coronavirus infection.’, Journal of molecular cell biology, 12(12), p. 968–979. doi: 10.1093/jmcb/mjaa042.

von Weyhern, C. H. et al. (2020) ‘Early evidence of pronounced brain involvement in fatal COVID-19 outcomes.’, Lancet (London, England), p. e109. doi: 10.1016/S0140-6736(20)31282-4.

Winkler, E. S. et al. (2020) ‘SARS-CoV-2 infection of human ACE2-transgenic mice causes severe lung inflammation and impaired function.’, Nature immunology. United States, 21(11), p. 1327–1335. doi: 10.1038/s41590-020-0778-2.

Xia, X., Wang, Y. and Zheng, J. (2021) ‘COVID-19 and Alzheimer’s disease: how one crisis worsens the other.’, Translational neurodegeneration. England, 10(1), p. 15. doi: 10.1186/s40035-021-00237-2.

Yin, K., Baillie, G. J. and Vetter, I. (2016) ‘Neuronal cell lines as model dorsal root ganglion neurons: A transcriptomic comparison.’, Molecular pain. United States, 12. doi: 10.1177/1744806916646111.

Yong, S. J. (2021) ‘Long COVID or post-COVID-19 syndrome: putative pathophysiology, risk factors, and treatments.’, Infectious diseases (London, England), 53(10), p. 737–754. doi: 10.1080/23744235.2021.1924397.

Yüce, M., Filiztekin, E. and Özkaya, K. G. (2021) ‘COVID-19 diagnosis -A review of current methods.’, Biosensors & bioelectronics, 172, p. 112752. doi: 10.1016/j.bios.2020.112752.

