## Supplementary figures and images for "SARS-CoV-2 infection leads to Tau pathological signature in neurons"

### Additional file 1.tiff

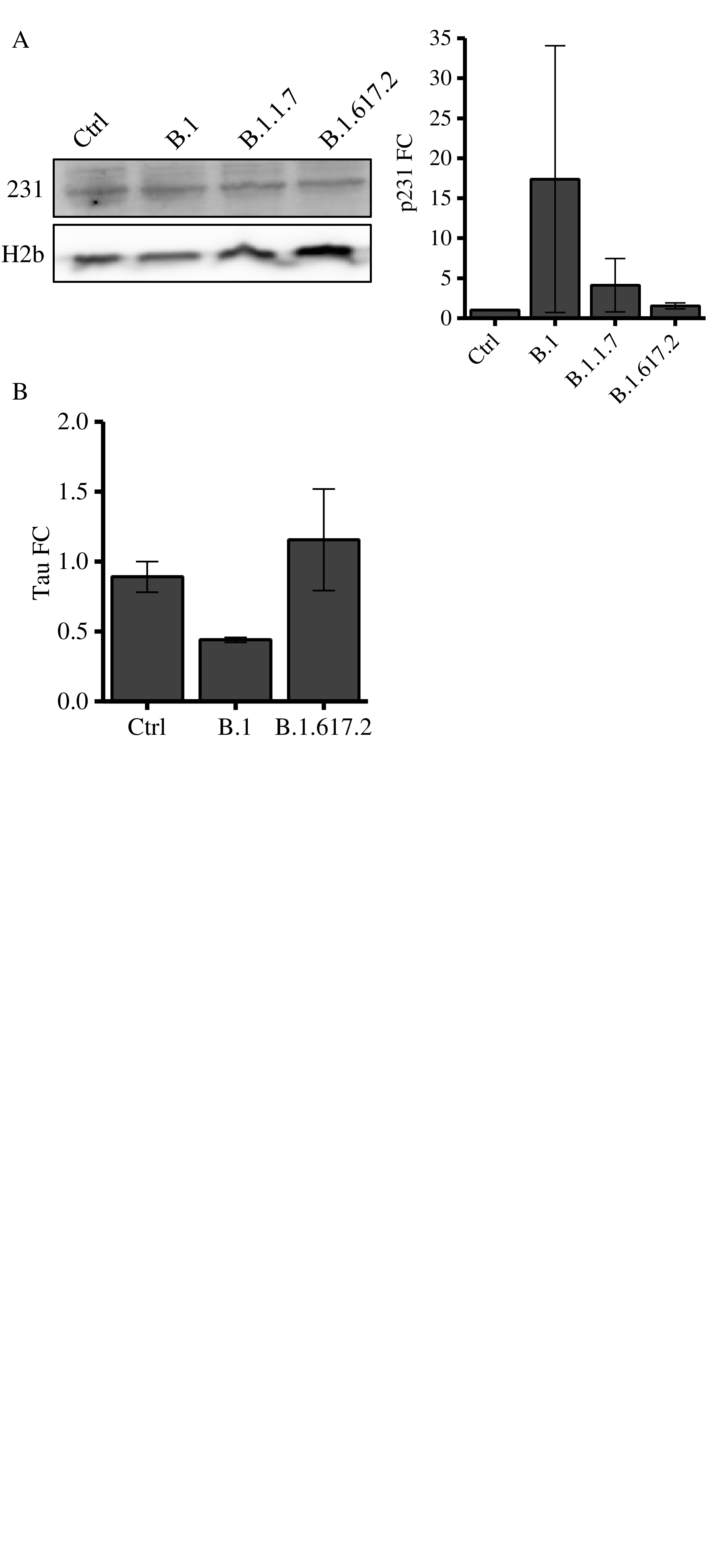

### Additional file 2.tiff

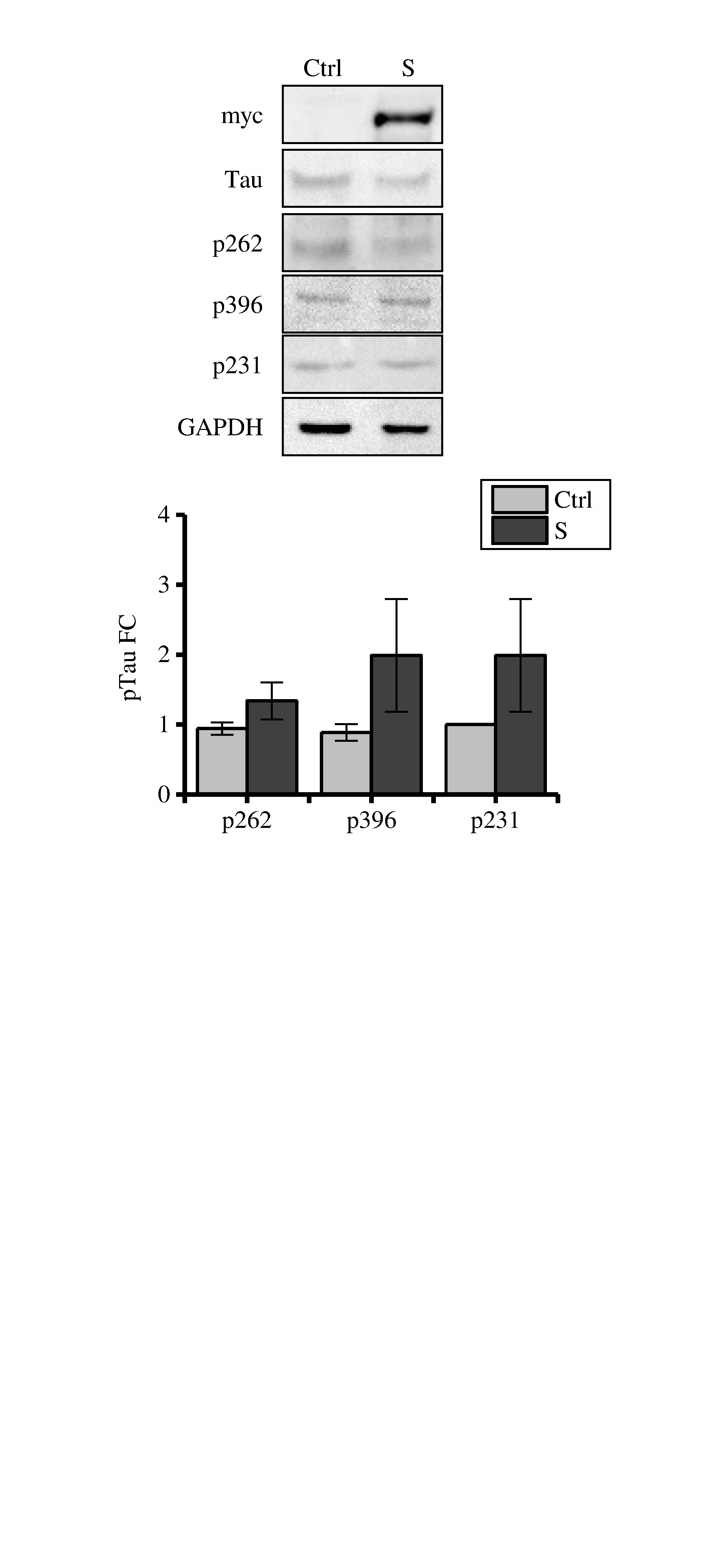
